# Characterization of the Golgi c10orf76-PI4KB complex, and its necessity for Golgi PI4P levels and enterovirus replication

**DOI:** 10.1101/634592

**Authors:** J.A. McPhail, H.R. Lyoo, J.G. Pemberton, R.M. Hoffmann, W. van Elst, J.R.P.M. Strating, M.L. Jenkins, J.T.B. Stariha, F.J.M. van Kuppeveld, T. Balla, J.E. Burke

**Affiliations:** Department of Biochemistry and Microbiology, University of Victoria, Victoria, BC, Canada; Department of Infectious Diseases & Immunology, Virology Division, Faculty of Veterinary Medicine, Utrecht University, Utrecht, The Netherlands; Section on Molecular Signal Transduction, Eunice Kennedy Shriver National Institute of Child Health and Human Development, National Institutes of Health, Bethesda, MD, USA

**Keywords:** c10orf76, ARMH3, phosphatidylinositol 4-kinase, PI4KB, PI4KIIIß, GBF1, Arf1, phosphoinositide, phosphatidylinositol 4-phosphate, PI4P, PKA, enterovirus, coxsackievirus A10, viral replication, hydrogen deuterium exchange mass spectrometry, HDX-MS, Golgi

## Abstract

The lipid kinase PI4KB, which generates phosphatidylinositol 4-phosphate (PI4P), is a key enzyme in regulating membrane transport and is also hijacked by multiple picornaviruses to mediate viral replication. PI4KB can interact with multiple protein binding partners, which are differentially manipulated by picornaviruses to facilitate replication. The protein c10orf76 is a PI4KB-associated protein that increases PI4P levels at the Golgi, and is essential for the viral replication of specific enteroviruses. We used hydrogen deuterium exchange mass spectrometry to characterize the c10orf76-PI4KB complex and reveal that binding is mediated by the kinase linker of PI4KB, with formation of the heterodimeric complex modulated by PKA-dependent phosphorylation. Complex-disrupting mutations demonstrate that PI4KB is required for membrane recruitment of c10orf76 to the Golgi, and that an intact c10orf76-PI4KB complex is required for the replication of c10orf76-dependent enteroviruses. Intriguingly, c10orf76 was also required for proper Arf1 activation at the Golgi, providing a putative mechanism for the c10orf76-dependent increase in PI4P levels at the Golgi.

**Highlights:**

- c10orf76 forms a direct complex with PI4KB, with the interface formed by a disorder-to-order transition in the kinase linker of PI4KB
- The c10orf76 binding site of PI4KB can be phosphorylated by PKA, with phosphorylation leading to decreased affinity for c10orf76
- Complex-disrupting mutants of PI4KB and c10orf76 reveal that PI4KB recruits c10orf76 to the Golgi/TGN
- Depletion of c10orf76 leads to decreases in both active Arf1 and Golgi PI4P levels
- Enteroviruses that rely on c10orf76 for replication depend on formation of the c10orf76-PI4KB complex

## Introduction

Phosphoinositides are essential regulatory lipids that play important roles in myriad cellular functions. The phosphoinositide species phosphatidylinositol-4-phosphate (PI4P) is widely distributed and involved in the coordinated regulation of membrane trafficking, cell division, and lipid transport [1,2]. Multiple human pathogens manipulate PI4P levels to mediate their intracellular replication, including *Legionella* [3] and multiple picornaviruses [4,5]. PI4P in humans is generated through the action of four distinct phosphatidylinositol-4-kinases: PI4KIIα (PI4K2A), PI4KIIβ (PI4K2B), PI4KIIIα (PI4KA) and PI4KIIIβ (PI4KB) [6-8]. PI4KB is localized at the Golgi and trans-Golgi-network (TGN), with PI4P pools in the Golgi apparatus generated by both PI4K2A and PI4KB [9]. While the localization and activity of PI4K2A is regulated through its palmitoylation, local membrane composition, and cholesterol levels [10], the activity of PI4KB is regulated by multiple protein-protein interactions [11-14]. These regulatory protein-protein interactions are in turn manipulated by many pathogenic RNA viruses that have evolved the ability to hijack PI4KB and generate PI4P-enriched replication organelles, which are essential for viral replication [15]. For most picornaviruses, manipulation of PI4P levels is driven by the action of the viral 3A protein and its interactions with PI4KB-binding proteins [13,16-18].

PI4KB plays both important catalytic and non-catalytic functions, with its regulation controlled by interactions with multiple protein binding partners, including acyl CoA Binding Domain 3 (ACBD3), Rab11, 14-3-3, and c10orf76 (chromosome 10, open-reading frame 76, also referred to as Armadillo-like helical domain-containing protein 3 (ARMH3)). PI4KB is a multi-domain lipid kinase containing a disordered N-terminus, a helical domain, and a bi-lobal kinase domain [14,19]. Biophysical and biochemical studies have defined the domains of PI4KB that mediate complex formation with a number of binding partners. The helical domain of PI4KB forms a non-canonical interaction with the small GTPase Rab11a that mediates localization of a pool of Rab11 to the Golgi and TGN [14,20]. PI4KB is primarily localized to the Golgi through the interaction of its N-terminus with ACBD3 [12,13]. PI4KB is activated downstream of ADP-ribosylation factor 1 (Arf1) [21], however, no evidence for a direct Arf1-PI4KB interface has been found, suggesting that this may be an indirect effect. PI4KB contains phosphorylation sites in disordered linkers between domains, including Ser294 in the helical-kinase linker of PI4KB, which is phosphorylated by protein kinase D (PKD). Phosphorylation of Ser294 drives binding of 14-3-3, which stabilizes PI4KB, prevents degradation, and increases Golgi PI4P levels [22-24]. Ser496 in the N-lobe linker of PI4KB is phosphorylated by protein kinase A (PKA) [25], and drives PI4KB localization from the Golgi to nuclear speckles [26]. c10orf76 was identified as a putative PI4KB interacting partner in immunoprecipitation experiments [17,27], with knockout of c10orf76 leading to decreased Golgi PI4P levels [28]. The function of this protein is unknown, however, it contains a domain of unknown function (DUF1741) that is well conserved in many eukaryotes.

Enterovirus proteins do not interact directly with PI4KB – they instead recruit PI4KB-regulatory proteins. A key component of manipulating PI4KB to generate PI4P-enriched replication organelles is the interaction of viral 3A proteins with host PI4KB-binding proteins. The 3A proteins from enteroviruses (i.e. Poliovirus, Rhinovirus, Coxsackievirus, Rhinovirus and Enterovirus 71) and Aichivirus recruit PI4KB to replication organelles through an interaction with ACBD3 [11,13,16-18,29-32]. The viral 3A protein from Aichivirus forms a direct interaction with the GOLD domain of ACBD3, leading to redistribution of PI4KB to replication organelles [11,31]. Enteroviruses also manipulate other lipid signaling pathways, with viral 3A proteins able to recruit the protein Golgi-specific brefeldin A-resistance guanine nucleotide exchange factor 1 (GBF1) that activates Arf1 [4,33-35], and subvert Rab11-positive recycling endosomes to replication organelles [36]. A new component of the PI4KB hijacking process, c10orf76, was identified as a key host factor in the replication of coxsackievirus A10 (CVA10) replication, but not coxsackievirus B1 (CVB1) [28].

We hypothesized that a direct c10orf76-PI4KB interaction may be critical for the regulation of Golgi PI4P levels and play a role in enterovirus replication. To elucidate the role of c10orf76 in PI4KB-mediated signaling, we utilized a synergy of hydrogen deuterium exchange mass spectrometry (HDX-MS) and biochemical assays to characterize the novel c10orf76-PI4KB complex *in vitro*. This allowed us to engineer complex-disrupting mutations that were subsequently used to define the role of the c10orf76-PI4KB complex in Golgi PI4P-signaling and viral replication *in vivo*. We find that PI4KB and c10orf76 form a high affinity complex mediated by a disorder-to-order transition of the kinase linker of PI4KB, with complex affinity modulated by PKA phosphorylation of the c10orf76 binding site on PI4KB. Knockout of c10orf76 lead to decreased PI4P levels, and disruption of Arf1 activation in cells. Complex-disrupting mutations revealed that c10orf76 is recruited to the Golgi by PI4KB, and that viral replication of enteroviruses that require c10orf76 is mediated by the c10orf76-PI4KB complex.

## Results

### c10orf76 forms a direct, high affinity complex with PI4KB

c10orf76 was previously identified as a putative PI4KB-binding partner through immunoprecipitation experiments [17,27], however, it was not clear if this was through a direct interaction. To identify a potential direct interaction between PI4KB and c10orf76 *in vitro*, we purified recombinant full-length proteins using a baculovirus and *Spodoptera frugiperda (Sf9)* expression system. Experiments on PI4KB used the slightly smaller isoform 2 variant (1-801, uniprot: Q9UBF8-2), compared to PI4KB isoform 1 (1-816 uniprot: Q9UBF8-1), similar to previous structural studies [14]. His-pulldown assays using NiNTA-agarose beads and purified recombinant proteins showed a direct interaction between PI4KB and His-tagged c10orf76 (**Fig. 1A**). Pulldown experiments carried out with the PI4KB-binding partners Rab11 and ACBD3 revealed that c10orf76 could form PI4KB-containing ternary complexes with both (**Fig. S1**), indicating a unique c10orf76 binding interface on PI4KB compared to Rab11 and ACBD3. To examine the stoichiometry of the c10orf76-PI4KB complex, we subjected apo c10orf76 and c10orf76 with PI4KB to size exclusion chromatography. Apo c10orf76 (79 kDa) eluted from the size exclusion column at a volume consistent with a monomer, while the c10orf76-PI4KB complex eluted at a volume consistent with a 1:1 complex (**Fig. 1B).** Since cellular knockout of c10orf76 has been shown to reduce PI4P levels *in vivo* [28], we investigated the effect of c10orf76 on PI4KB lipid kinase activity with biochemical membrane reconstitution assays using phosphatidylinositol (PI) vesicles. Intriguingly, c10orf76 was a potent inhibitor of PI4KB, with inhibition being dose-dependent and possessing an IC_50_ of ∼90 nM (**Fig. 1C**). This inhibitory effect was observed on both pure phosphatidylinositol (PI) vesicles, and vesicles that mimic the composition of the Golgi (20% PI, 10% PS, 45% PE, 25% PC) (**Fig. 1D**). This paradoxical PI4KB-inhibitory result *in vitro* conflicts with observed Golgi PI4P decreases in c10orf76 deficient cells [28]. This suggests that biochemical assays may not fully recapitulate the environment of the Golgi. To further define the role of this complex we focused on defining the molecular basis of this interface, allowing for generation of binding-deficient mutants for downstream cellular and viral experiments.

**Figure 1.**
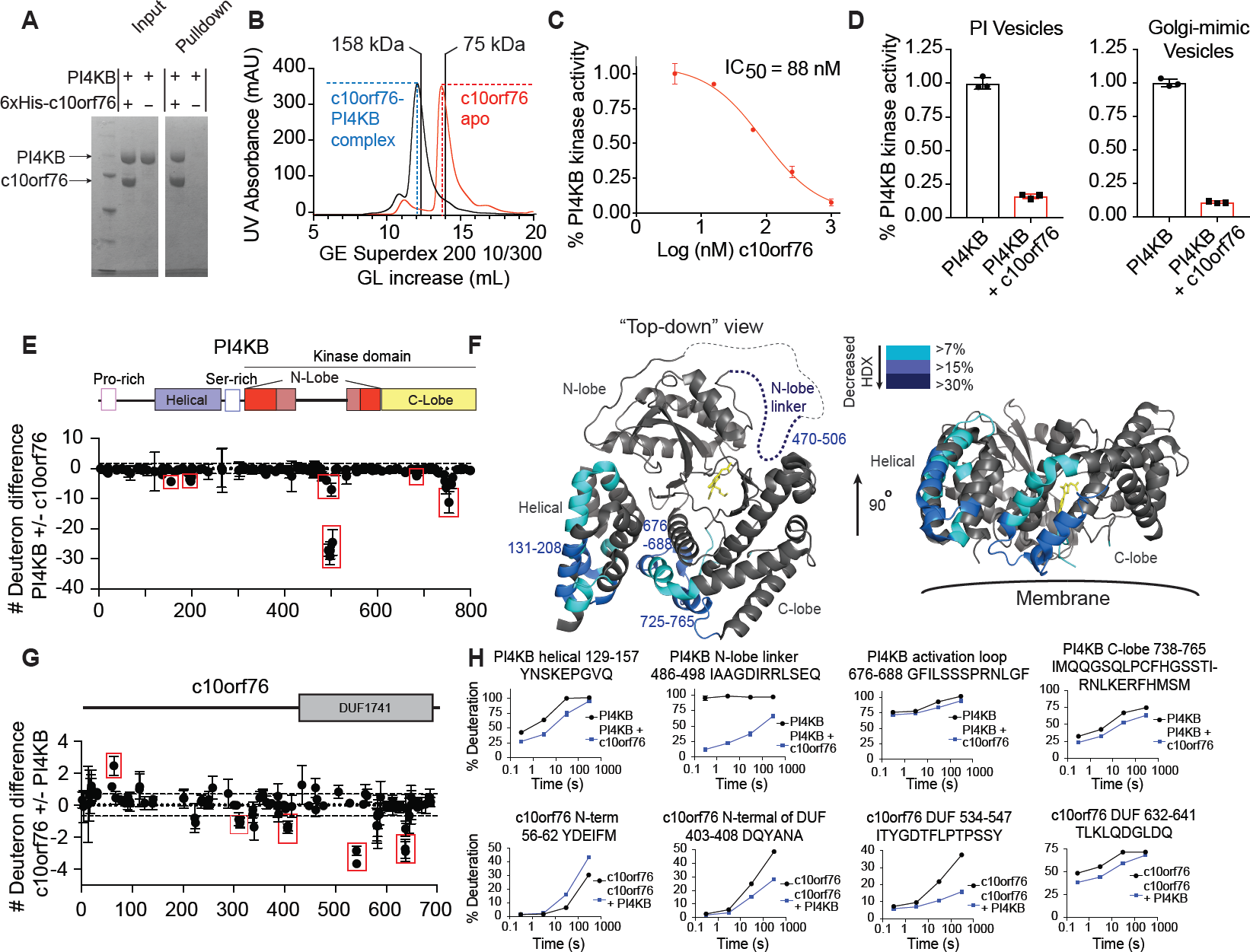
c10orf76 directly binds to PI4KB through an extended interface focused at the N-lobe kinase linker of PI4KB. **(A)** Recombinant c10orf76 directly binds to PI4KB *in vitro.* His-pulldown assays of baculovirus/*Sf9* produced 6xHis-c10orf76 (3 μM) were carried out with untagged PI4KB (2.5 μM). **(B)** PI4KB and c10orf76 form a stable complex. The complex of c10orf76-PI4KB eluted from a S200 superdex 10/300 GL increase gel filtration column (GE) at a volume consistent with a heterodimer (169 kDa), while c10orf76 alone eluted at a volume consistent with a monomer (79 kDa). Lines with MW values indicate elution of MW standards (158 kDa aldolase, 75 kDa conalbumin). **(C)** PI4KB is potently inhibited by c10orf76 in a dose-dependent manner *in vitro*. Kinase assays of PI4KB (20 nM) in the presence of varying concentrations of c10orf76 (1.6 nM-1 μM) were carried out on pure PI lipid vesicles (0.5 mg/L) in the presence of 100 μM ATP. The data was normalized to the kinase activity of PI4KB alone. IC_50_ values were determined by one binding site, nonlinear regression (curve fit) using Graphpad. Error bars represent standard deviation (n=3). **(D)** PI4KB is potently inhibited by c10orf76 on pure PI vesicles and vesicles mimicking Golgi composition. Kinase assays of PI4KB and c10orf76 were carried out on lipid substrate composed of pure PI vesicles (0.5 mg/mL) with 100 μM ATP, and Golgi mimic vesicles (0.5 mg/ml, 10% PS, 20% PI, 25% PE, 45% PC) with 10 μM ATP. PI4KB was present at 20 and 300 nM in the PI and Golgi substrate assays respectively, with c10orf76 present at 500 nM in both experiments. The data is normalized to the kinase activity of PI4KB alone. Error bars represent standard deviation (n=3). **(E)** Changes in deuterium incorporation PI4KB in the presence of c10orf76 showed a profound ordering of the kinase domain N-lobe linker and smaller changes in the helical domain and C-lobe of the kinase domain. The sum of the difference mapped as the difference in number of deuterons incorporated for PI4KB (400 nM) in the presence and absence of c10orf76 (400 nM) over all time points (3s at 1 °C; 3s, 30s, and 300s at 23 °C). Each dot represents a peptide graphed on the x-axis according to the central residue. The red boxes highlight key regions that showed significant changes (>7% decrease in exchange, >0.5 Da difference, and unpaired two-tailed student t-test p<0.05). For all panels error bars represent standard deviation (n=3). **(F)** c10orf76 binding induces differences in HDX throughout multiple domains of PI4KB. Regions of >7% difference in deuterium exchange in the presence of c10orf76 are colored onto the structure of PI4KB according to the legend (PDB: 4D0L). The N-lobe linker of the kinase domain is disordered in the structure and is represented by a dotted line. **(G)** Changes in the deuterium incorporation of c10orf76 in the presence of PI4KB. H/D exchange reactions displayed as the sum of the difference in HDX in the number of deuterons for c10orf76 (400 nM) in the presence of PI4KB (400 nM) at all time points (3s at 1 °C; 3s, 30s, and 300s at 23 °C) analyzed. Red boxes highlight regions that showed significant changes (>7% decrease in exchange, >0.5 Da difference, and unpaired two-tailed student t-test p<0.05). **(H)** The PI4KB N-lobe linker undergoes a disorder-to-order transition upon binding c10orf76. Selected peptides (including the sequence, domain information, and numbering) of both PI4KB and c10orf76 displayed as the % deuteration incorporation over time.

### HDX-MS reveals that PI4KB and c10orf76 form an extended interface involving a disorder-to-order transition of the PI4KB N-lobe linker

To identify the putative interface between PI4KB and c10orf76, we employed hydrogen-deuterium exchange mass spectrometry (HDX-MS) to map regions protected in both proteins upon complex formation. HDX-MS is an analytical technique that measures the exchange rate of amide hydrogens in proteins. Because one of the main determinants for amide exchange is the presence of secondary structure, their exchange rate is an excellent readout of protein dynamics. HDX-MS is thus a potent tool to determine protein-protein, protein-ligand, and protein-membrane interactions [37-39]. H/D exchange was carried out for three different conditions: apo PI4KB, apo c10orf76, and a 1:1 complex of PI4KB with c10orf76. Deuterium incorporation experiments were carried out at four different timepoints (3, 30 and 300 seconds at 23°C and 3 seconds at 1°C). Deuterium incorporation is determined by quenching the exchange reaction in a solution that dramatically decreases the exchange rate, followed by rapid digestion, peptide separation, and mass analysis. A total of 185 peptides covering 96.9% of the PI4KB sequence, and 108 peptides covering 73.9% of the c10orf76 sequence were generated (**Fig. 1E-H-source data 1, Fig. S2**). Significant differences in deuterium exchange between conditions were defined as changes in exchange at any timepoint that met the three following criteria: greater than 7% change in deuterium incorporation, a greater than 0.5 Da difference in peptide mass, and a p-value of less than 0.05 (unpaired student’s t-test).

Multiple regions of PI4KB were protected from amide exchange in the presence of c10orf76, revealing an extended binding interface (**Fig. 1E,F,H; Fig. S2)**. The most prominent difference in exchange was at the C-terminus of the disordered N-lobe linker (residues 486-498), where the presence of c10orf76 led to a significant ordering of this region. This region had no protection from amide exchange in the apo state, revealing it to be disordered, with a very strong stabilization (>80% decrease in exchange) in the presence of c10orf76, indicating a disorder-to-order transition upon c10orf76 binding (**Fig. 1H**). This N-lobe kinase linker is dispensable for lipid kinase activity, as it can be removed with a minimal effect on PI4KB catalytic activity [19]. In addition to this change there were multiple smaller decreases in exchange in the helical domain (131-138, 149-157, 159-164, and 183-204) and kinase domain (676-688, 725-734, and 738-765). The helical domain of PI4KB mediates binding to Rab11. However, the PI4KB-Rab11 complex was still able to form in the presence of c10orf76 (**Fig. S1)**. The decreases in exchange with c10orf76 observed in the kinase domain were located in the activation loop (676-688) and the C-lobe (738-765), which may mediate the inhibition observed *in vitro*. The protected surface on PI4KB extensively spans the membrane face of the kinase, which may prevent PI4KB from directly interfacing with the membrane and accessing PI in the presence of c10orf76, at least in the absence of other binding partners *in vitro* (**Fig. 1F).**

The presence of PI4KB also caused multiple differences in H/D exchange in c10orf76, with increased exchange at the N-terminus (56-62) as well as decreased exchange N-terminal of, and within, the domain of unknown function (DUF1741; 403-408, 534-547, 632-641) (**Fig. 1G,H; Fig. S2**). There are no clear structural determinants of c10orf76, with limited homology to any previously solved structure, however, it is predicted to consist of a primarily helical fold arranged into armadillo repeats. The uncharacterized DUF1741 domain of c10orf76 is present throughout many eukaryotes, however, even though the DUF1741 domain is strongly conserved in evolution, c10orf76 is the only protein that contains this domain in humans.

The largest observed change in deuterium incorporation in either protein was in the PI4KB N-lobe linker (486-496). Interestingly, this region contains a consensus PKA motif (RRxS) that corresponds to Ser496 (Ser511 in PI4KB isoform 1), which is phosphorylated *in vivo* and conserved back to the teleost fishes (**Fig. 2A**) [26]. Systems level analysis of PKA signaling networks also show that phosphorylation of this site is decreased >90% in PKA knockout cells, indicating that it is likely a direct PKA target [25]. To better understand the regulation of the c10orf76-PI4KB complex, we sought to characterize the effects of Ser496 phosphorylation.

**Figure 2.**
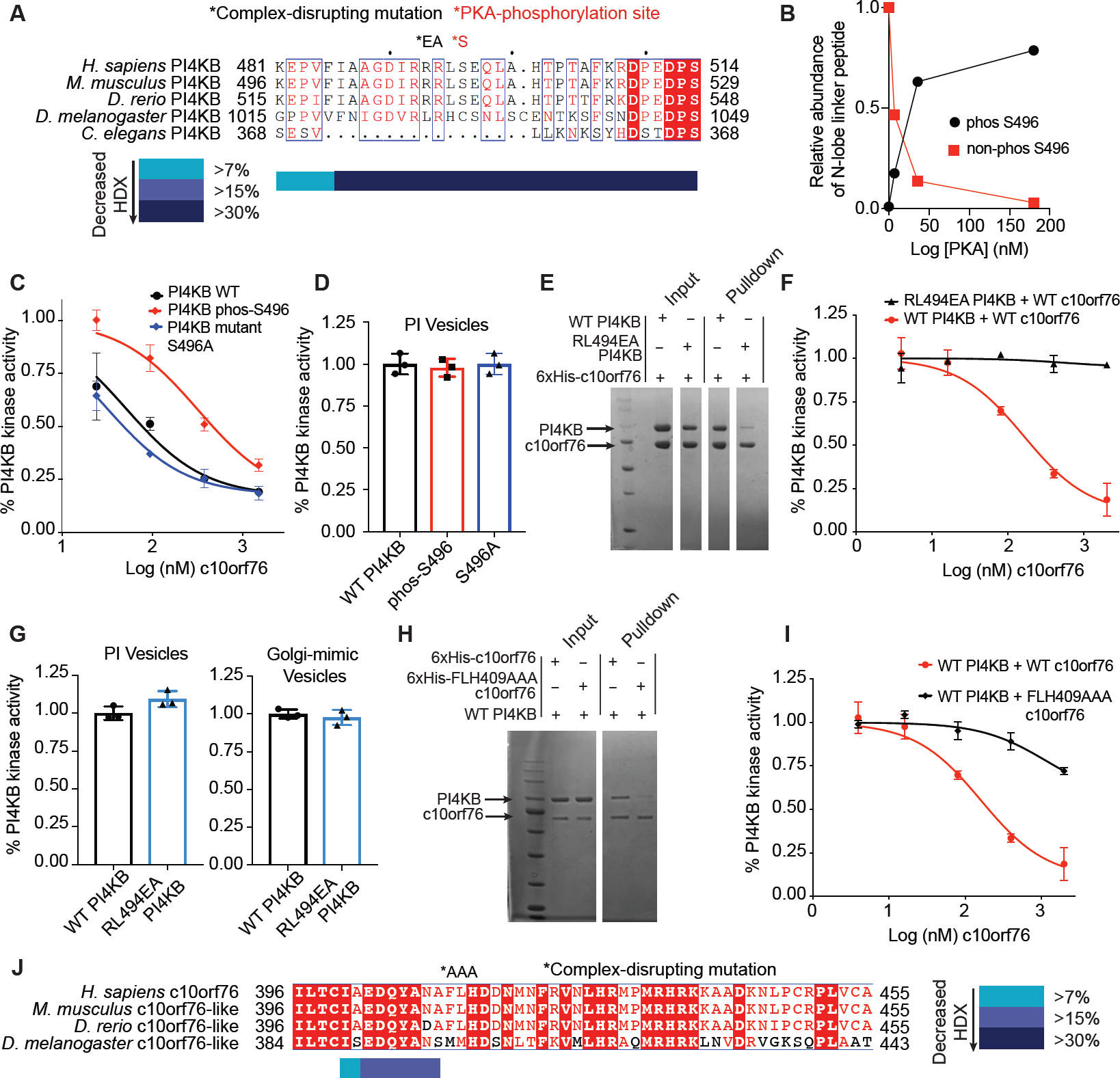
The PI4KB c10orf76 interface is conserved and can be post-translationally modified by PKA, with rationally designed mutations disrupting complex formation. **(A)** The N-lobe kinase linker region of PI4KB is strongly conserved back to *D. rerio*. The N-lobe linker region of PI4KB sequences of the organisms indicated were analysed using Clustal Omega/ ESpript 3. The consensus PKA motif (RRXS) that is conserved back to *D. rerio* is indicated on the sequence, as well as the RL494EA point mutation. **(B)** The N-lobe kinase linker of PI4KB can be efficiently phosphorylated by PKA. Recombinant PKA at different concentrations (0, 7, 34, 168, or 840 ng) was incubated with recombinant (*E. coli*) wild-type PI4KB (20 μg) for 1 hour with 200 μM ATP and the amount of phosphorylation was followed using mass spectrometry. Relative abundance of Ser496 phosphorylated PI4KB was calculated using the relative intensity (total area) of the phosphorylated vs non-phosphorylated peptide (486-506). **(C)** PI4KB phosphorylation by PKA alters the affinity for c10orf76. The kinase activity of different variants of PI4KB (15 nM) was measured in the presence of varying amounts of c10orf76 (1.6 nM-2μM) with 100% PI lipid substrate (0.5 mg/L) and 100 μM ATP. The data was normalized to the kinase activity of PI4KB alone Error bars represent standard deviation (n=3). **(D)** PI4KB has the same kinase activity when Ser496 is phosphorylated or mutated to alanine. Kinase assay of PI4KB non-phosphorylated, phos-Ser496 or S496A (15 nM) on pure PI lipid vesicles (0.5 mg/L) with 100 μM ATP. The data was normalized to the kinase activity of WT PI4KB. Error bars represent standard deviation (n=3). **(E)** Engineered RL494EA PI4KB mutant shows decreased binding to c10orf76. His-pulldown assays of 6xHis-c10orf76 (3 μM) with wild-type or RL494EA PI4KB (1-2 μM). **(F)** RL494EA PI4KB activity is not inhibited by c10orf76. Kinase assays of either wild type or mutant RL494EA PI4KB (40 nM) were carried out with varying concentrations of c10orf76 (3.9 nM-2μM) with 100% PI lipid vesicles (0.5 mg/L) and 100 μM ATP. The data was normalized to the kinase activity of PI4KB alone. Error bars represent standard deviation (n=3). **(G)** Wild-type PI4KB and RL494EA PI4KB mutant have the same lipid kinase activity. Kinase assays of either wild-type and mutant PI4KB (10 nM) were carried out with 100% PI lipid vesicles (0.5 mg/L), 100 μM ATP, and PI4KB (300 nM) on Golgi-mimic vesicles (0.5 mg/mL) with 10 μM ATP. The data was normalized to the kinase activity of WT PI4KB. Error bars represent standard deviation (n=3). **(H)** FLH409AAA-c10orf76 mutant shows decreased affinity for PI4KB. His-pulldown assays of 6xHis-c10orf76 (1 μM) with wild-type PI4KB (1 μM). Samples washed a total of 4 times. **(I)** Kinase assay shows FLH409AAA c10orf76 inhibition of PI4KB is greatly reduced. Kinase assay of PI4KB (40 nM) and a concentration curve of c10orf76 (3.9 nM-2μM) on pure PI lipid vesicles (0.5 mg/L) with 100 μM ATP. The data was normalized to the kinase activity of PI4KB alone. Error bars represent standard deviation (n=3). **(J)** The PI4KB-binding region of c10orf76 is strongly conserved back to *D. rerio*. Clustal Omega/ ESpript 3 alignment of the FLH409 region of c10orf76 that binds PI4KB.

### PI4KB is directly phosphorylated at Ser496 by PKA to modulate the affinity of the c10orf76-PI4KB complex

There are three well-validated phosphorylation sites on PI4KB: Ser294, Ser413, Ser496 [40]. To test the role of phosphorylation of PI4KB at Ser496, we generated stoichiometrically phosphorylated PI4KB at only Ser496 using an *in vitro* phosphorylation approach that relied on the production of the purified mouse PKA catalytic subunit in *E. coli*. To minimize complications from any background phosphorylation that occurs in *Sf9* cells, we used PI4KB expressed in *E. coli* to ensure the starting protein substrate was non-phosphorylated. Analysis of the *Sf9*-produced PI4KB revealed significant phosphorylation of Ser294, Ser413, Ser430 and Ser496, while *Sf9*-produced c10orf76 had evidence of phosphorylation of Ser14, and an additional Ser/Thr phosphorylation in the 325-351 region, although the specific residue is ambiguous from the MS data. No phosphorylation was identified from *E. coli* produced proteins, as expected (**Fig. S3**). Dose response assays for the phosphorylation of PI4KB Ser496 using *E. coli*-produced protein were then carried out with increasing concentrations of purified PKA, and the resulting product was analyzed by mass spectrometry for the site-specific incorporation of the phosphate moiety (**Fig. 2B)**. Ser496 in PI4KB was phosphorylated efficiently by PKA, with >99% phosphorylation at Ser496 occurring with a 1:500 ratio of PKA to PI4KB and no detectable phosphorylation at the other major PI4KB phosphorylation sites (**Fig. S3**). Lipid kinase assays were then carried out using different concentrations of c10orf76 for both phosphorylated and non-phosphorylated PI4KB. The phosphorylated form had a 3-fold increase in the IC_50_ value, suggesting that Ser496 phosphorylation decreases c10orf76 binding affinity, with no shift in the IC_50_ value for the S496A PI4KB mutant (**Fig. 2C)**. Kinase assays carried out on both Ser496 phosphorylated PI4KB and non-phosphorylated PI4KB showed that there is no direct effect of the phosphorylation events on basal lipid kinase activity (**Fig. 2D**). PKA-mediated phosphorylation-dependent changes in the affinity of protein-protein complexes have been previously described [41,42]. We utilized HDX-MS to test if the altered inhibition profile we saw was due to decreased affinity between c10orf76 and PI4KB. These experiments were carried out at a single time point of D_2_O exposure (5 seconds at 20°C) with differing levels of c10orf76 present. Plotting the difference in deuterium incorporation versus c10orf76 concentration gives a characteristic binding isotherm for both phosphorylated and non-phosphorylated PI4KB; displaying a ∼3-fold decreased affinity for the phosphorylated form of PI4KB (85 nM vs 30 nM, **Fig. S3)** Phosphomimic variants of Ser496 in PI4KB mutants did not alter the affinity for c10orf76, so they could not be utilized to study this effect *in vivo* (data not shown). To better characterize the role of the c10orf76-PI4KB complex *in vivo*, we sought to generate c10orf76-PI4KB complex-disrupting mutations.

### Rationally engineered PI4KB and c10orf76 mutants that disrupt complex formation

The c10orf76 binding site within the N-lobe kinase linker of PI4KB identified by HDX-MS is highly conserved in vertebrates, with much of the region also conserved in *D. melanogaster*, but not in *C. elegans* (**Fig. 2A**). We used a combination of both the sequence conservation and HDX-MS results to design a complex-disrupting mutant. The RL residues at 494-495 were mutated to EA (RL494EA), effectively causing both a charge reversal and decrease in hydrophobicity. The RL494EA mutant disrupted binding to His-tagged c10orf76 bait in a His pulldown assay (**Fig. 2E**) and prevented inhibition by c10orf76 in kinase assays (**Fig. 2F**). This mutant had exactly the same basal kinase activity as the WT PI4KB on both PI vesicles and Golgi-mimetic vesicles (**Fig. 2G)**, strongly suggesting that the mutant kinase is properly folded. In an attempt to design rational mutations of c10orf76 that also disrupted binding to PI4KB, multiple mutations were tested in regions 403-408, 534-547 and 632-641 that were identified using HDX-MS. Combining the HDX-MS data and sequence homology, we designed a triple alanine mutant at the end of a putative helix (QYANAFL) that was well conserved in vertebrates (**Fig. 2J)**, close to the HDX-MS protection (FLH residues 409-411 to AAA, referred to as FLH mutant afterwards). The FLH mutant expressed well, significantly reduced binding to PI4KB in a His-pulldown assay (**Fig. 2E**), and also showed a marked reduction in its ability to inhibit PI4KB activity (**Fig. 2I**). To confirm the c10orf76 FLH mutant does not affect global protein structure, we compared deuterium incorporation of the c10orf76 wild-type and FLH mutant and observed no changes in deuterium incorporation seen outside of the predicted helix containing the FLH residues (**Fig. S4).** The engineering of complex-disrupting mutants that do not alter catalytic activity or protein folding provided an excellent tool to test the importance of the c10orf76-PI4KB complex in cells.

### PI4KB recruits c10orf76 to the Golgi

To define the role of the c10orf76-PI4KB interface in cellular localization we utilized fluorescently-tagged variants of the wild-type and complex-disrupting mutants of both PI4KB and c10orf76. Fluorescence microscopy of HEK293 cells expressing GFP-tagged wild-type PI4KB revealed that it primarily localizes to the Golgi (**Fig. 3A**). GFP-PI4KB RL494EA, which is deficient in c10orf76 binding, also localized mainly to the Golgi, which suggests that c10orf76 plays a minimal role in the Golgi recruitment of PI4KB (**Fig. 3A**). The wild-type GFP-c10orf76 also localizes to the Golgi. However, the PI4KB binding-deficient FLH mutant is redistributed to the cytosol; revealing an important role for PI4KB in the proper cellular localization of c10orf76 (**Fig. 3B**). To further analyze the role of PI4KB in the recruitment of c10orf76, we utilized a chemically-inducible protein heterodimerization system that relies on the selective interaction of the FKBP12 (FK506 binding protein 12) and FRB (a 9 kDa fragment of mTOR that binds rapamycin) modules upon treatment with rapamycin [12,43]. Specifically, we fused the FRB domain to residues 34–63 of a CFP-tagged mitochondrial localization signal from mitochondrial A-kinase anchor protein 1 (AKAP1), and fused mRFP-FKBP12 onto the wild-type or mutant variants of human PI4KB (**Fig. 3C**). These constructs allowed us to examine the localization of the wild-type or mutant GFP-c10orf76 following the acute sequestration of PI4KB to the outer mitochondrial membrane, where other Golgi-associating proteins are not be present. Treatment with rapamycin (100 nM) caused the rapid recruitment of mRFP-FKBP12-PI4KB to the mitochondria, which also caused the rapid co-recruitment of c10orf76 (**Fig. 3D; Video 1**); suggesting that PI4KB is the only component necessary for membrane recruitment of c10orf76. Experiments using mRFP-FKBP12 PI4KB RL494EA showed that although the mutant kinase is relocated to the mitochondria, GFP-c10orf76 does not co-localize (**Fig. 3E; Fig. 3F, Video 2**). Taken together, these live-cell studies corroborated the protein interaction studies completed *in vitro* and also demonstrate that the newly defined c10orf76-PI4KB interface is required for proper localization of c10orf76 to the Golgi. Compellingly, these findings reveal a potential novel function of PI4KB in the recruitment of c10orf76.

**Figure 3.**
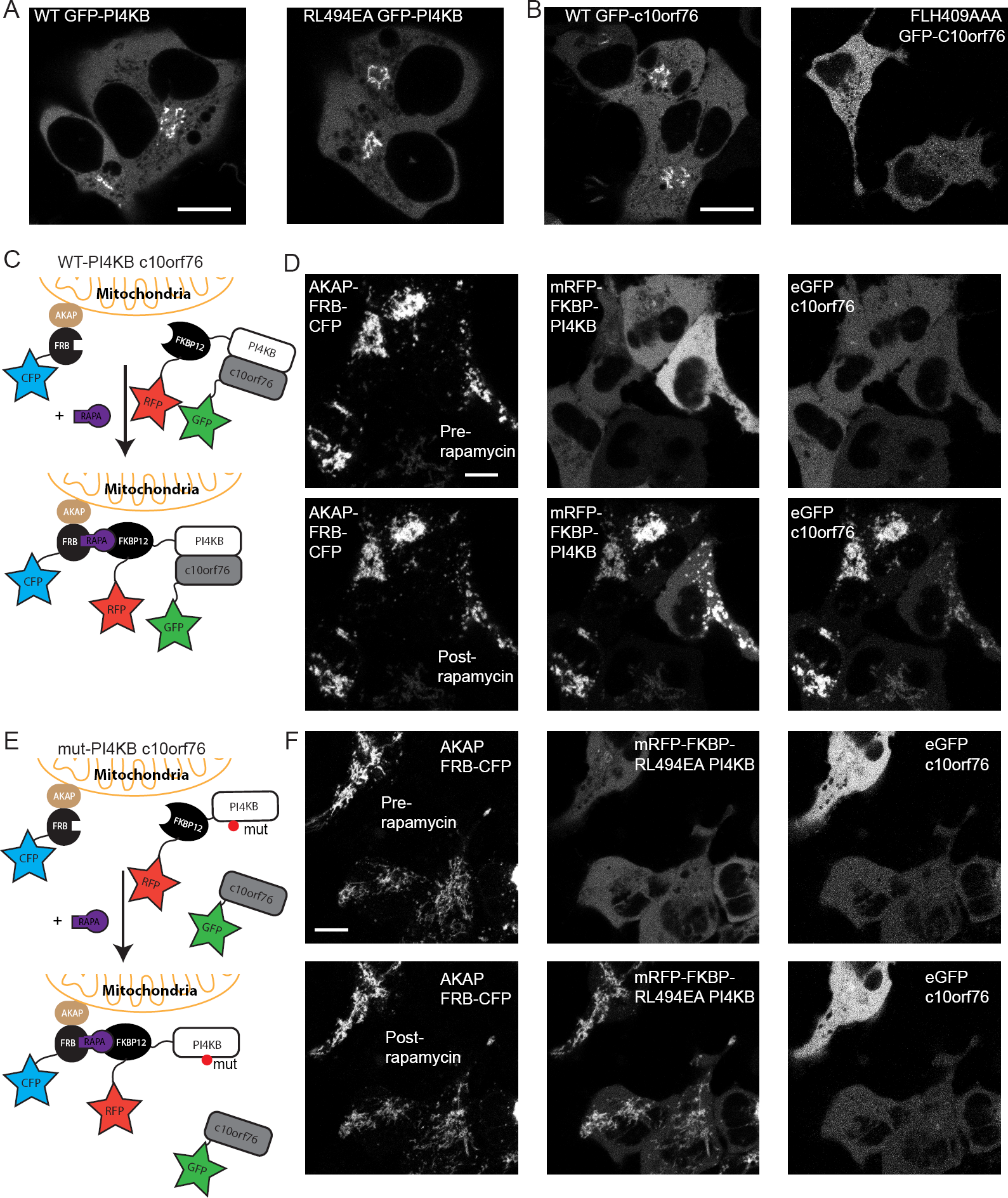
PI4KB recruits c10orf76 to the Golgi *in vivo*. **(A)** Transfections of HEK293 cells revealed that both wild-type GFP-PI4KB and RL494EA GFP-PI4KB localize to the Golgi. **(B)** WT c10orf76 also localized to the Golgi, however, the PI4KB binding deficient mutant of c10orf76 (FLH409AAA) predominantly localized to the cytosol. **(C)** Cartoon schematic of rapamycin-inducible mitochondria recruitment. The AKAP1-FRB-CFP construct is localized to the outer mitochondrial membrane, while the RFP-FKBP12-PI4KB and GFP-c10orf76 are localized in the Golgi as well as within the cytoplasm where they can form a complex. Upon addition of rapamycin, the RFP-FKBP12-PI4KB construct is translocated to the mitochondria. **(D)** Mitochondria recruitment experiment with wild-type PI4KB and c10orf76. Left: AKAP1-FRB-CFP is localized to the mitochondria before (top) and 5 minutes after rapamycin (100 nM) treatment (bottom). Middle: RFP-FKBP12-PI4KB is located in the cytosol before rapamycin (top) and translocates to the mitochondria after rapamycin induction (bottom). Right: GFP-c10orf76 is located in the cytosol before rapamycin (top) and translocates to the mitochondria after rapamycin induction (bottom). **(E)** Schematic of the rapamycin-inducible mitochondria recruitment experiment with mutant PI4KB and WT c10orf76. **(F)** Mitochondria recruitment experiment with mutant PI4KB and WT c10orf76. Left: AKAP1-FRB-CFP is localized to the mitochondria before (top) and 5 minutes after (bottom) rapamycin treatment. Middle: RFP-FKBP12-PI4KB(RL494EA) is located in the cytosol before rapamycin (top) and translocates to the mitochondria after rapamycin induction (bottom). Right: GFP-c10orf76 is located in the cytosol before (top) and after (bottom) rapamycin induction. Bars represent 10 μm.

### c10orf76 regulates Arf1 activation and maintains Golgi PI4P levels

The paradoxical finding that the loss of c10orf76 leads to increased PI4P levels in cells, yet decreased catalytic activity of PI4KB *in vitro*, suggested that there was an unknown lipid or protein constituent in cells that is not present in our *in vitro* experiments. To determine the role of c10orf76 in cells we examined the distribution of different Golgi-localized signaling components in c10orf76-deficient (knockout) HAP1 cells. In agreement with previous studies [28], we found that there were decreased PI4P levels at the Golgi in c10orf76 knockout cells, as indicated by decreased Golgi staining by an anti-PI4P antibody (**Fig. 4A**). Intriguingly, there was an apparent increase in Golgi localized PI4KB in the c10orf76 knockout cells (**Fig. 4A**), similar to what occurs upon treatment with a PI4KB inhibitor [44], clearly indicating that decreased PI4P production was not due to loss of PI4KB recruitment in the absence of c10orf76. We tested the localization of different Golgi markers to verify that decreased PI4P was not due to disruption of Golgi morphology. Markers for the cis Golgi (GM130), cis/medial Golgi (Giantin), and the trans-Golgi network (TGN46) all showed similar localization in both WT and c10orf76 knockout cells (**Fig. 4B**). The distribution of the ER-Golgi intermediate compartment marker ERGIC53 was also similar, suggesting that Golgi morphology was maintained in the c10orf76 knockout HAP1 cells (**Fig. 4B**).

**Figure 4.**
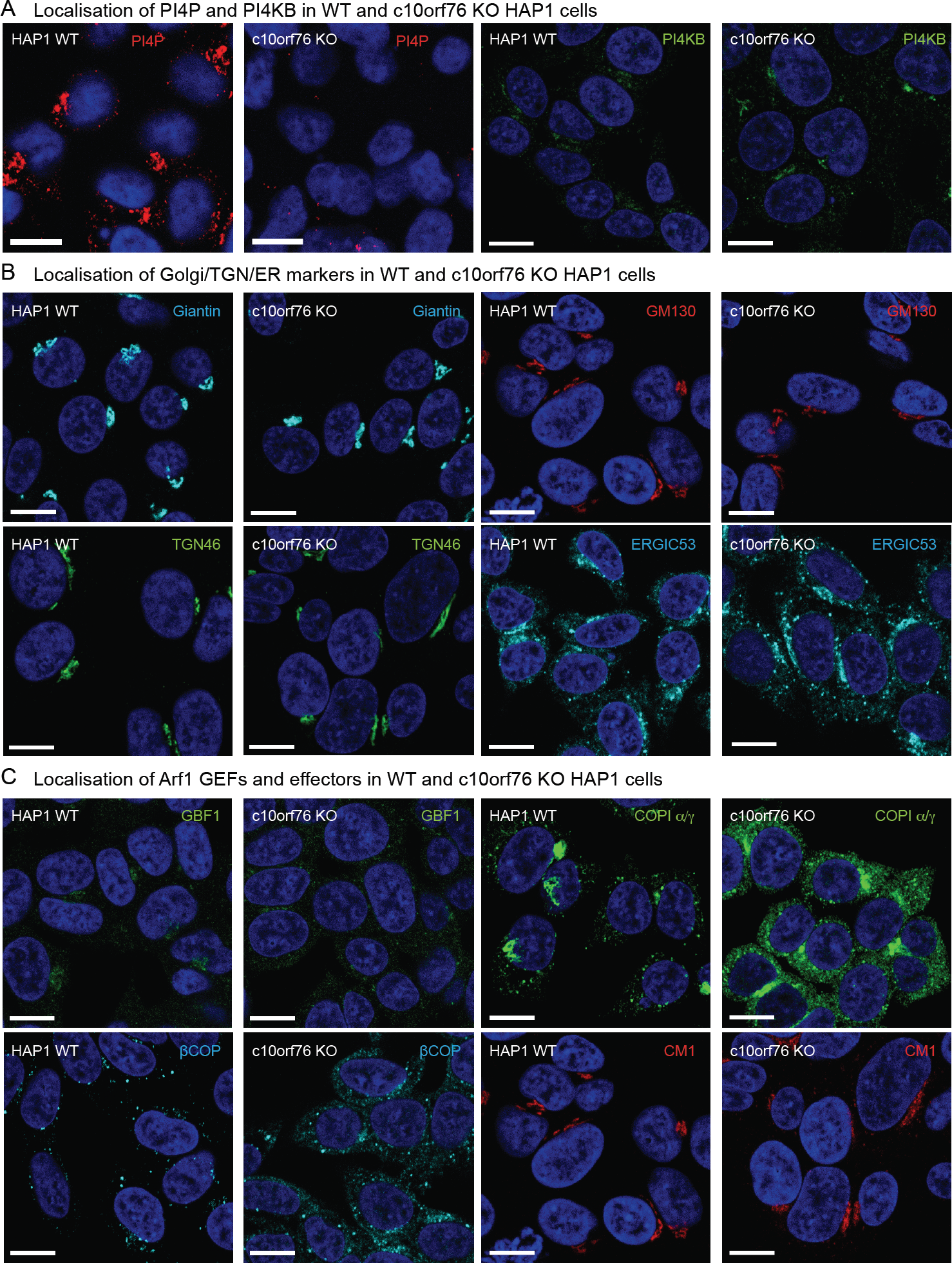
Knockout of c10orf76 in HAP1 cells leads to decreased PI4P levels and disruption of GBF1 / active Arf1 localization despite minimal effects on Golgi morphology. HAP1 cells were fixed and stained with antibodies examining PI4P and PI4KB (**A**), Golgi morphology markers (**B**), and markers of Arf1 activation (**C**). The coatomer proteins COPIα/γ and βCOP act as a readout for GTP-bound Arf1, while the native coatomer was detected with the CM1 antibody. Nuclei were stained with DAPI (blue). Bars represent 10 μm.

We next tested the localization of the Arf1-GEF GBF1, as active GTP-bound Arf1 is a putative activator of PI4KB [21]. In c10orf76 knockout cells there was a redistribution of GBF1, with GBF1 being more diffuse, with less localized at the Golgi (**Fig. 4C**). The generation of active GTP-bound Arf1 by Arf-GEFs leads to recruitment of multiple effector proteins, with one of most well characterised being the coatomer proteins, which form COPI coated vesicles that mediate Golgi to ER trafficking. Antibody staining with the CM1 antibody, which only recognizes the native form of coatomer, showed similar Golgi distribution for both WT and c10orf76 knockout cells. However, antibodies recognizing COP-β and -α/γ subunits, which associate with GTP-bound Arf1, not only showed staining in the Golgi, but also diffuse staining in the cytosol in c10orf76 knockout cells which was not observed in wild type cells (**Fig. 4C**). Together, these results suggest that c10orf76 plays a key role in Arf1 activation, likely providing a mechanism for increased PI4P levels driven by c10orf76.

### Replication of c10orf76-dependent enteroviruses requires intact c10orf76-PI4KB interaction

All enteroviruses depend on PI4KB kinase activity for replication. Despite the physical and functional connection between PI4KB and c10orf76, enteroviruses showed different dependencies on c10orf76 [28]. Specifically, while Coxsackievirus A10 (CVA10) replication was impaired in c10orf76 knockout cells, the replication of CVB1 was not. Furthermore, c10orf76 was identified as a pro-viral factor for replication of poliovirus (PV1) [45]. We set out to investigate the importance of the c10orf76-PI4KB interaction for replication of CVA10 and PV1. We first made a side-by-side comparison of virus replication in HAP1 wildtype and c10orf76 knockout cells in a single cycle of replication. The replication of CVA10 was significantly impaired in c10orf76 deficient cells, with partial inhibition of PV1 replication, and no impairment for replication of CVB3 (**Fig. 5A**). Due to the notoriously difficult nature of transfecting HAP1 cells, we determined the importance of the c10orf76-PI4KB interaction for virus replication in HeLa PI4KB knockout cells transfected with different PI4KB expression plasmids as previously described [29]. Expression of wild type PI4KB efficiently restored the replication of all viruses (**Fig. 5B**). Expression of the PI4KB RL494EA mutant that is deficient in binding c10orf76 fully rescued replication in CVB3, only partially rescued PV1 replication, and failed to rescue CVA10 replication. These observations suggest that the c10orf76-PI4KB interaction is necessary for CVA10, and to a lesser extent, PV1 replication and thereby implies that functions of c10orf76 are selectively hijacked by specific viruses.

**Figure 5.**
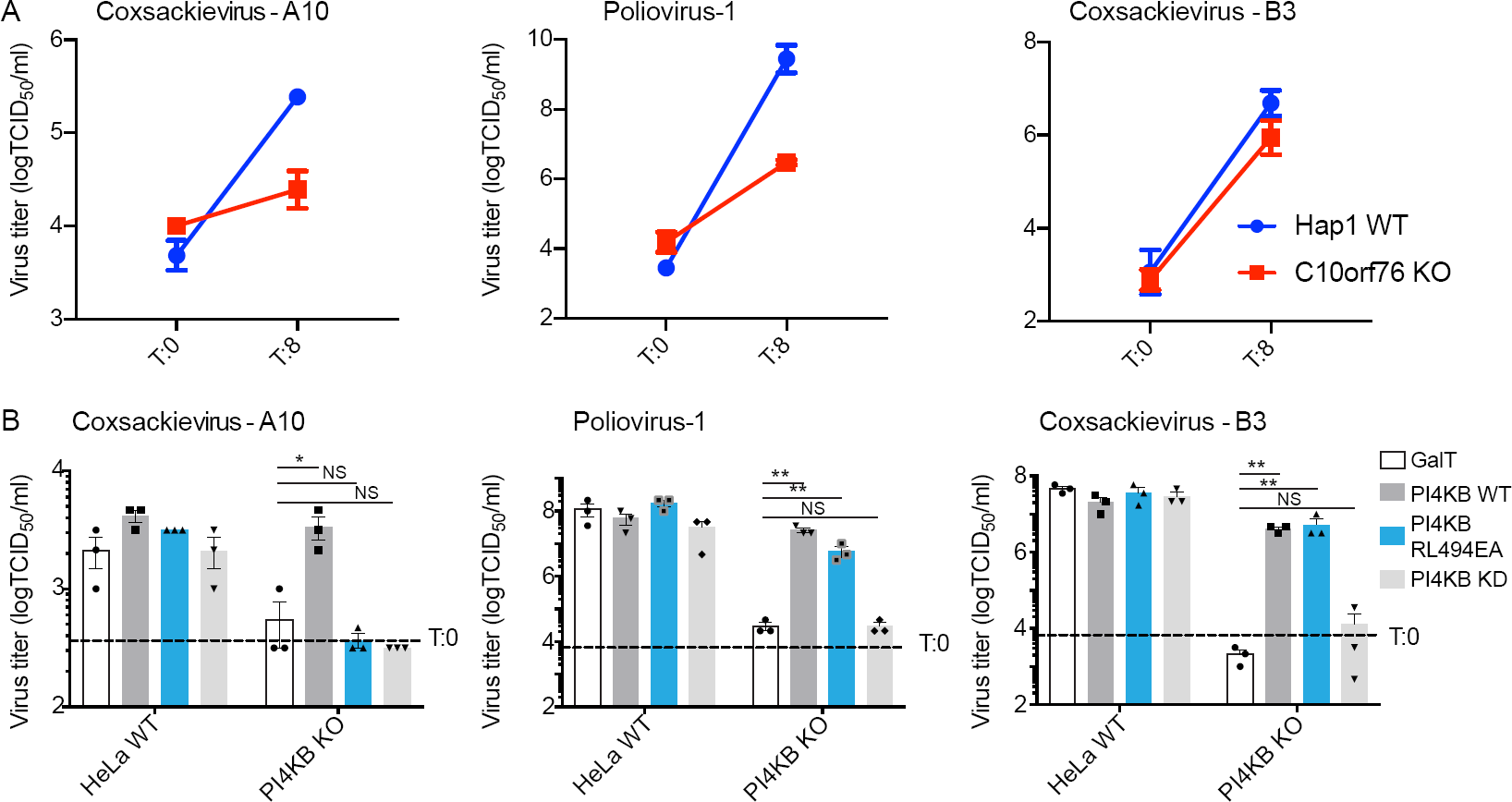
The c10orf76-PI4KB complex is essential for Coxsackievirus A10 replication. **(A)** Viral infection assays determining viral titers by end-point titration at 0 hours and 8 hours in HAP1 wild-type or c10orf76 knockout cells. Left: Coxsackievirus A10 infection. Middle: Poliovirus-1 infection. Right: Coxsackievirus B3 infection. **(B)** Viral infection assays determining virus titers by end point titration at 8 hours in HeLa wild-type and PI4KB knockout cells upon transfection of wild-type PI4KB, the complex-disrupting RL494EA PI4KB mutant or the kinase dead D674A PI4KB mutant. Left: Coxsackievirus A10 infection. Middle: Poliovirus-1 infection. Right: Coxsackievirus B3 infection. Values were statistically evaluated compared to the GalT control using a one-way ANOVA. **, P<0.01; *, P<0.05; N.S., P>0.05. For all panels error bars represent standard error (n=3).

## Discussion

Defining the full complement of cellular roles for PI4KB is an important objective in characterizing the integrated control of secretion and membrane trafficking at the Golgi, and also provides a framework for understanding how PI4P can be manipulated by viruses. We have identified the c10orf76-PI4KB interaction as an important Golgi signaling complex and a critical factor in the replication of specific enteroviruses. Multiple mechanisms have been previously described for how PI4KB participates in Golgi signaling and membrane trafficking, including detailed insights into protein binding partners, post-translational modifications, and regulated recruitment to specific membrane compartments. PI4KB was originally identified in yeast (yeast protein PIK1) as an essential gene [46], with its activity playing a key role in secretion from the Golgi [47]. The mammalian isoform was identified soon afterwards through its sensitivity to wortmannin [48-50]. The first identified Golgi activator of PI4KB was the GTPase Arf1 [21]. However, no direct interaction has been established, which indicates a potential indirect mechanism of activation. Phosphorylation of PI4KB by PKD at Ser294 mediates binding to 14-3-3 proteins, with this leading to an increase in PI4KB activity [22,23], that has been suggested to correspond with an increase in PI4KB stability [24]. The most well validated protein binding partner that regulates Golgi recruitment of PI4KB is ACBD3 (previously referred to as GCP60) [13]. ACBD3 forms a direct, high-affinity interface with PI4KB that is mediated by a disorder-to-order transition in the N-terminus of PI4KB upon binding to the Q domain of ACBD3 [11,12]. The recruitment of PI4KB to the Golgi by ACBD3 is controlled through the direct interaction of the GOLD domain of ACBD3 with the Golgi resident transmembrane protein Giantin [51]. In addition to regulatory protein interactions, PI4KB is predicted to contain an ALPS motif at the C-terminus that mediates lipid binding to unsaturated membranes [9]. PI4KB plays key non-catalytic roles through its interaction with the GTPase Rab11, with PI4KB required for localizing a pool of Rab11 to the Golgi and TGN [52]. This interaction is mediated through a non-canonical, nucleotide-independent binding interface with the helical domain of PI4KB [14]. However, there are still many unexplained aspects of PI4KB recruitment and regulation, highlighted by the increased recruitment of PI4KB to the Golgi following treatment with PI4KB inhibitors that is concomitant with a decrease in Golgi PI4P levels [44].

The protein c10orf76 was originally identified as a putative PI4KB interacting partner through co-immunoprecipitation experiments using tagged PI4KB [17,27]. Tests of genetic essentiality identified c10orf76 as a central molecular hub at the Golgi, with it being synthetically lethal in combination with the loss of several different Golgi-signaling proteins, and also showing a genetic link to PI4KB [28]. That study also found that c10orf76 is essential in the KBM7 CML cell line, but not in HAP1 cells, with this relationship also being true for PI4KB. Additional evidence on the essentiality of this protein is highlighted by the homozygous mutant of ARMH3, the mouse homolog of c10orf76, which is lethal at the pre-weaning stage [53]. c10orf76 is highly conserved in vertebrates and we find a strong correlation between the conservation of the kinase linker region of PI4KB and the PI4KB-binding site in c10orf76, suggesting that a key role of c10orf76 is linked to its ability to form a complex with PI4KB. PI4KB recruitment to the Golgi is not mediated by c10orf76, but instead it appears that PI4KB is responsible for the Golgi-recruitment of c10orf76. *In vitro*, c10orf76 led to decreased lipid kinase activity of PI4KB. However, knockout of c10orf76 in cells led to reduced PI4P levels. This discrepancy could be due to the lack of other interacting partners *in vitro*, such as Arf1/GBF1. c10orf76 knockout led to an increased cytosolic fraction of Arf1 effectors and the Arf GEF GBF1. Our work reveals c10orf76 as a novel player in Arf1 regulation, with c10orf76 required for maintaining active Arf1 and corresponding Golgi PI4P levels.

Enteroviruses hijack numerous lipid signaling processes within infected cells to mediate their replication through the generation of replication organelles, with recruitment of PI4KB [4] and GBF1 [33] playing key roles in this process. Recruitment of these cellular host factors in enteroviruses is primarily mediated through the action of membrane-bound viral 3A proteins, which form either direct or indirect interactions that are important for facilitating replication organelle formation. One of the most well-conserved 3A binding partners in enteroviruses is the Golgi resident protein ACBD3, which interacts with the central part of 3A and recruits as well as activates PI4KB [11,13,18,29,31]. The N-terminal part of the 3A proteins from several enteroviruses (*e.g.*, poliovirus and coxsackie virus B3) directly binds and recruits GBF1, but this interaction is less conserved, severely reduced, or even absent in the 3A proteins of rhinoviruses due to subtle amino acid differences in their N-terminus [18,33]. We find that c10orf76 is required for replication of coxsackie virus A10 and, to a lesser extent, poliovirus and that c10orf76-dependent viruses rely on the c10orf76-PI4KB interface. Poliovirus is the causative agent of poliomyelitis, and coxsackie virus A10 is an important cause of outbreaks of hand-foot-and-mouth disease, but which is also associated with severe, and sometime fatal, clinical symptoms such as aseptic meningitis. Remarkably, replication of coxsackie virus B1 [28] and coxsackie virus B3 (this study) is independent of c10orf76. Why the c10orf76-PI4KB interface is necessary for replication of some enteroviruses, but not others, is unknown. The differential dependence on c10orf76 could possibly be explained by distinct affinity of 3A proteins from different viruses towards GBF1. Alternatively, each virus may require specific threshold PI4P level for efficient formation of its replication complexes or replication organelles. More research on the dependence of viral replication on either GBF1 or c10orf76-mediated alteration of PI4P levels is required to better understand how enteroviruses hijack these complex membrane trafficking processes.

Direct inhibition of PI4KB is likely not a useful antiviral strategy due to unexpected deleterious side effects of PI4KB inhibition in animal models [54]. The targeting of other cellular host factors used to manipulate PI4KB signaling or feedback is a potential avenue for development of novel antiviral therapeutics. Identification of a direct high-affinity c10orf76-PI4KB complex that regulates the cellular localization of c10orf76 represents key insight into the multifaceted regulation of PI4KB signaling. The important role of the c10orf76-PI4KB complex in the replication of select enteroviruses represents a novel molecular platform which is targeted by viruses that hijack lipid signaling. The involvement of c10orf76 in Arf1 dynamics, as well as the dependence on PI4KB for Golgi localization of c10orf76, reveals a potential role of the c10orf76-PI4KB complex in Arf1 activation and subsequent PI4P production.

## Materials and Methods

### Protein expression and purification

#### c10orf76 and PI4KB

The human *C10orf76* gene (Uniprot Q5T2E6) was synthesized by GeneArt (Thermofisher). c10orf76 and PI4KB (Uniprot Q9UBF8-2) were each expressed with an N-terminal 6xHis-tag followed by a TEV protease site. The c10orf76 and PI4KB proteins purified for HDX-MS were expressed in *Spodoptera frugiperda* (*Sf9*) cells by infecting 1-4 L of cells at a density of 1.5 x 10^6^ cells/mL with baculovirus encoding the kinase. After 60-72 hours infection at 27°C, *Sf9* cells were harvested and washed in phosphate-buffered saline (PBS). The c10orf76 and PI4KB proteins utilized for assays, mutational analysis and studying PKA phosphorylation were expressed in Rosetta (DE3) *E. coli* (c10orf76) or BL21 C-41(DE3) *E. coli* (PI4KB) induced overnight at 16 °C with 0.1 mM IPTG at an OD_600_ of 0.6. Cell pellets containing c10orf76 or PI4KB were sonicated in NiNTA Buffer (20 mM Tris-HCl pH 8.0, 100 mM NaCl, 20 mM imidazole, 5% (v/v) glycerol, 2 mM β-mercaptoethanol) containing protease inhibitors (Millipore Protease Inhibitor Cocktail Set III, Animal-Free) for 5 minutes on ice. Triton X-100 (0.1% v/v) was added to the cell lysate and the lysed cell solution was centrifuged for 45 minutes at 20,000 × g at 2°C. Supernatant was filtered through a 5 μm filter and loaded onto a 5 mL HisTrap™ FF crude (GE) column in NiNTA buffer. The column was washed with 1.0 M NaCl and 20 mM imidazole in NiNTA buffer and protein was eluted with 200-250 mM imidazole in NiNTA buffer. Eluted c10orf76 or PI4KB was pooled and concentrated onto a 5 mL HiTrap™ Q column (GE) equilibrated with Q buffer (20 mM Tris-HCl pH 8.0, 100 mM NaCl, 5% glycerol v/v, 2 mM β-mercaptoethanol) and eluted with an increasing concentration of NaCl. Protein was pooled and concentrated using an Amicon 30K concentrator and incubated overnight on ice with the addition of TEV protease. Size exclusion chromatography (SEC) was performed using a Superdex™ 200 10/300 GL increase (GE) column equilibrated in SEC buffer (20 mM HEPES pH 7.5, 150 mM NaCl and 0.5 mM TCEP). Fractions containing the protein of interest were pooled, concentrated, spun down to remove potential aggregate and flash frozen in liquid nitrogen for storage at −80 °C. c10orf76-PI4KB complex SEC trace was generated by mixing c10orf76 and PI4KB in a 1:1 ratio after individual anion exchange runs and then injecting onto the Superdex™ 200 10/300 GL increase (GE) column. Elution volumes of protein standards were obtained from the GE Instruction 29027271 AH Size exclusion chromatography columns document. See *Protein Kinase A (PKA) treatment of PI4KB* for details on producing the phosphorylated variant of PI4KB.

#### ACBD3, Rab11a and PKA

ACBD3 and Rab11a were expressed with N-terminal GST tags, with Protein kinase A (*M. musculus* PKA catalytic subunit alpha; Addgene 14921) expressed with an N-terminal His tag. ACBD3, Rab11a, and PKA were expressed in BL21 C-41(DE3) *E. coli* cells, with ACBD3 and Rab11 expression carried out overnight at 16 °C with 0.1 mM IPTG, and PKA expression was carried out for 4 hours at 28 °C with 1 mM IPTG. ACBD3, Rab11, and PKA were purified as previously published [11,14,55]. In brief, cell pellets containing expressed ACBD3 or Rab11a were sonicated in Q Buffer (20 mM Tris-HCl pH 8.0, 100 mM NaCl, 5% (v/v) glycerol, 2 mM β-mercaptoethanol) containing protease inhibitors (Millipore Protease Inhibitor Cocktail Set III, Animal-Free) for 5 minutes on ice. Triton X-100 (0.1% v/v) was added to the cell lysate and the lysed cell solution was centrifuged for 45 minutes at 20,000 x g at 2°C. Supernatant was filtered through a 5 μm filter and incubated with 1-4mL of Glutathione Sepharose™ 4B beads (GE) for 1-2 hours at 4°C. Beads were then washed with Q buffer, and GST-tagged proteins were eluted with 20 mM glutathione in Q buffer. Protein was further purified using anion exchange and size-exclusion chromatography as described above and final protein was spun down to remove potential aggregate and flash frozen in liquid nitrogen for storage at −80 °C. Nickel purification of PKA proceeded as described for PI4KB, and nickel elute was concentrated, spun down to remove potential aggregate and flash frozen in liquid nitrogen for storage at −80 °C.

### Nickel and GST Pulldown Assays

For His pulldowns, NiNTA agarose beads (Qiagen) (20 μL) were washed three times by centrifugation and resuspension in NiNTA buffer. His-tagged bait protein was then added to a concentration of 1-3 µM and incubated with the beads on ice for 10 minutes in a total volume of 50 μL. Beads were washed three times with 150 μL NiNTA buffer at 4 °C. Non-His-tagged prey protein was then added to a final concentration of 1-2 µM in a total volume of 50 μL, at which point 10 μL was taken for SDS-PAGE analysis. The mixture was incubated on ice for an additional 30 minutes and then washed four times with 120 μL NiNTA buffer at 4 °C at which time an aliquot was taken as the output for SDS-PAGE analysis.

For GST pulldowns, Glutathione Sepharose™ 4B beads (GE healthcare) were washed three times by centrifugation and resuspension in Q buffer. GST-tagged bait protein (or control GST) was then added to a concentration of 3-6 µM in 50 μL and incubated with the beads on ice for 10 minutes in a total volume of 50 μL. Beads were washed three times with 150 μL Q buffer at 4 °C. Non-GST-tagged prey protein were then added to a final concentration of 2-4 µM in a total volume of 50 μL, at which point the input was taken for SDS-PAGE analysis. The mixture was incubated on ice for an additional 30 minutes and then washed four times with 120 μL Q buffer at 4 °C, at which time an aliquot was taken as the output for SDS-PAGE analysis.

### Vesicle Preparation and Lipid Kinase Assays

Lipid kinase assays were carried out using the Transcreener® ADP^2^ FI Assay (BellBrook Labs) following the published protocol as previously described [11]. In brief, substrate stocks were made up containing 1.0 mg/mL PI vesicles or 4.0 mg/mL Golgi-mimetic vesicles (10% PS, 20% PI, 25% PE, 45% PC) and were extruded through a 100 nm Nanosizer Extruder (T&T Scientific) and then combined with in a buffer containing 20 mM Hepes pH 7.5, 100 mM KCl and 0.5 mM EDTA (200 µM ATP with 1.0 mg/mL PI vesicles, 20 µM ATP with 1.0 mg/mL Golgi-mimetic vesicles). Kinase reactions were started by adding 2 µL of this substrate stock in a 384-well black low volume plates (Corning 4514). Proteins were thawed on ice and spun down to remove precipitate. Proteins were diluted individually to 4X the desired concentration in Kinase Buffer (40 mM Hepes pH 7.5, 200 mM NaCl, 20 mM MgCl_2_, 0.8% Triton-X, and 0.2 mM TCEP) on ice. Proteins were then mixed together or with additional Kinase buffer resulting in 2X desired concentrations of each protein. To start the reaction, 2 µL of 2X protein stock was added to 2 µL of 2X substrate stock in plates. After mixing, the 4 µL reactions consisted of 30 mM HEPES pH 7.5 (RT), 100 mM NaCl, 50 mM KCl, 10mM MgCl_2_, 0.25 mM EDTA, 0.4% (v/v) Triton-X, 0.1 mM TCEP, 10 µM ATP and 0.5 mg/mL vesicles. PI4KB was run at a final concentration of 15 nM, 20 nM or 40 nM and c10orf76 was run in 4-fold curves from 1 µM – 3.9 nM or 5-fold curves from 2 µM – 1.6 nM. Reactions proceeded at 23°C for 20-30 minutes. Reactions were stopped using 4 µL of the transcreener stop buffer (1X Stop & Detect Buffer B, 8 nM ADP Alexa594 Tracer, 97 µg/ml ADP2 Antibody-IRDye® QC-1). Fluorescence intensity was measured using a Spectramax M5 plate reader with λex x = 590 nm and λem m = 620 nm (20nm bandwidth). Data was plotted using Graphpad Prism software, with IC_50_ values determined by nonlinear regression (curve fit). No detectable nonspecific ATPase activity was detected in reactions containing 250 nM wild-type PI4KB without vesicle substrate.

### Mapping the c10orf76-PI4KB binding interface using HDX-MS

HDX reactions were conducted in 50 µL reactions with a final concentration of 400 nM of protein per sample (c10orf76-PI4KB, 400 nM each). Reactions were initiated by the addition of 45 µL of D_2_O Buffer Solution (10 mM HEPES pH 7.5, 50 mM NaCl, 97% D_2_O) to 5 µL of protein solution, to give a final concentration of 87% D_2_O. Exchange was carried out for four timepoints, (3s at 1°C and 3s, 30s, and 300s at 23 °C). Exchange was terminated by the addition of acidic quench buffer giving a final concentration 0.6 M guanidine-HCl and 0.8% formic acid. All experiments were carried out in triplicate. Samples were immediately frozen in liquid nitrogen and stored at −80°C until mass analysis.

### Comparison of FLH409AAA and WT c10orf76 secondary structure

HDX-MS reactions were performed with 40 µL final volume with a protein concentration of 0.25 µM in each sample. Reactions were started by the addition of 39 µL D2O buffer (100mM NaCl, 35 mM Hepes, 91.7% D_2_O) to 1 µL of protein (Final: 89.4% D2O). Reactions were quenched by the addition of 30uL of acidic quench buffer (3% formic acid, 2M Guanidine) resulting in final 1.28% Formic acid and 0.85M guandine-HCl. Proteins were allowed to undergo exchange reactions for either 3s or 300s at 23°C prior to addition of quench buffer and flash freezing in liquid N_2_. All samples were set and run in triplicate. Samples were stored at −80°C until injection onto the UPLC for MS analysis.

### HDX-MS data analysis

Protein samples were rapidly thawed and injected onto a UPLC system kept in a cold box at 2°C. The protein was run over two immobilized pepsin columns (Applied Biosystems; porosyme, 2-3131-00) stored at 10°C and 2°C at 200 µL/min for 3 min and the peptides were collected onto a VanGuard precolumn trap (Waters). The trap was subsequently eluted in line with an Acquity 1.7 µm particle, 100 × 1 mm^2^ C18 UPLC column (Waters), using a gradient of 5-36% B (buffer A 0.1% formic acid, buffer B 100% acetonitrile) over 16 minutes. MS experiments were performed on an Impact QTOF (Bruker) and peptide identification was done by running tandem MS (MS/MS) experiments run in data-dependent acquisition mode. The resulting MS/MS datasets were analyzed using PEAKS7 (PEAKS) and a false discovery rate was set at 1% using a database of purified proteins and known contaminants. HD-Examiner Software (Sierra Analytics) was used to automatically calculate the level of deuterium incorporation into each peptide. All peptides were manually inspected for correct charge state and presence of overlapping peptides. Deuteration levels were calculated using the centroid of the experimental isotope clusters. Attempts at generating fully deuterated protein samples to allow for the control of peptide back exchange levels during digestion and separation were made for all proteins. Protein was incubated with 3M guanidine for 30 minutes prior to the addition of D_2_O, where they were further incubated for an hour on ice. The reactions were then quenched as before. Generation of a fully deuterated sample was successful for PI4K using this method, however generation of fully deuterated c10orf76 failed. Results for c10orf76 are therefore presented as relative levels of deuterium incorporation and the only control for back exchange was the level of deuterium present in the buffer (87%). The average error of all time points and conditions for each HDX project was less than 0.2 Da. Therefore, changes in any peptide at any time point greater than both 7% and 0.5 Da between conditions with an unpaired t-test value of p<0.05 was considered significant. The full details of H/D exchange for all peptides are shown in Source data, with statistics described in **Supplemental Table 1**.

### Protein Kinase A (PKA) Treatment of PI4KB

PKA (mouse catalytic subunit) was serially diluted and different concentrations were incubated with PI4KB in 20 μL reactions on ice for 1 hour (20 μg PI4KB, 20 mM MgCl_2_, 200 μM ATP and either 840 ng, 168 ng, 34 ng, 7 ng or 0 ng PKA). Reactions were terminated by the addition of acidic quench buffer giving a final concentration 0.6 M guanidine-HCl and 0.8% formic acid and then flash frozen in liquid N_2_ prior to MS phosphorylation analysis.

To generate *E. coli* expressed, PKA phosphorylated PI4KB for use in kinase assays and HDX-MS, phosphorylation of Ser496 was carried out using 1.0 mg PI4KB, 20 mM MgCl_2_, 200 μM ATP and 4.2 μg PKA in NiNTA buffer, with the reaction allowed to proceed for 1 hour on ice. The reaction was quenched with 20 mM EDTA, and immediately loaded onto a GE 1 mL HisTap FF crude to remove His-tagged PKA. Phosphorylated PI4KB was concentrated followed by size exclusion chromatography as described for PI4KB above. In tandem, a non-phosphorylated PI4KB control was purified in the same manner except MgCl_2_, ATP, and PKA were not added. Protein was flash frozen in liquid N_2_ for storage at −80 °C.

### HDX-MS dose response of c10orf76 of phosphorylated PI4KB

Phosphorylated and non-phosphorylated PI4KB were generated and purified as described above. HDX reactions were conducted in 130 µl reaction volumes with a final concentration of 20nM PI4KB (phosphorylated or non-phosphorylated) per sample, with 0 nM, 5 nM, 10 nM, 20 nM, 40 nM, 80 nM, 160 nM and 320 nM c10orf76. Exchange was carried out for 5 seconds, in triplicate for each concentration of c10orf76. Hydrogen deuterium exchange was initiated by the addition of 80 µl of D_2_O buffer solution (10 mM HEPES (pH 7.5), 50 mM NaCl, 97% D_2_O) to the protein solution, to give a final concentration of 60% D_2_O. Exchange was terminated by the addition of 20 µl ice cold acidic quench buffer at a final concentration 0.6 M guanidine-HCl and 0.9% formic acid. Samples were immediately frozen in liquid nitrogen at − 80 °C. Data analyzed as described above in *HDX-MS data analysis.*

### Phosphorylation Analysis

LC-MS/MS analysis of phosphorylated variants of PI4KB was carried out as described in the HDX-MS data analysis section. MS/MS datasets were analyzed using PEAKS7 to identify phosphorylated peptides in PI4KB and c10orf76. A false discovery rate was set at 0.1% using a database of purified proteins and known contaminants. To measure PI4KB phosphorylation levels using Bruker Data analysis, the phosphorylated and non-phosphorylated peptides of interest were extracted, and the total area of each peptide was manually integrated to determine the amount of phosphorylated vs non-phosphorylated species under given experimental conditions. No phosphorylation was detected in *E. coli* derived PI4KB. For *Sf9* derived PI4KB Ser294 phosphorylation, the peptides KRTAS*NPKVENEDE (290-303) and KRTAS*NPKVENEDEPVRLADERE (290-312) were averaged, for Ser413 phosphorylation DTTSVPARIPENRIRSTRS*VENLPECGITHE (395-425) was used, for Ser430 phosphorylation GITHEQRAGS*F (430-441) was used, and for Ser496 phosphorylation IAAGDIRRRLS*EQLAHTPTA (486-505) and IAAGDIRRRLS*EQ-LAHTPTAF (486-506) were averaged. No phosphorylation was detected in *E. coli* derived c10orf76. For *Sf9* derived c10orf76 Ser14 phosphorylation, LRKSS*ASKKPLKE (10-22) was used, and for the 325-351 phosphorylation (exact location of phosphorylation ambiguous) VTTPVSPAPTTPVTPLGTTPPSSD (326-348), VTTPVSPAPTTPVTPLGTTPPSSDVISS (325-351) and VTTPVSPAPTTPVTPLGTTPPSS (325-347) were averaged.

### Alignments

Protein sequences from the Uniprot database were aligned using Clustal Omega [56] and figures were generated using ESPript [57]. Uniprot PI4KB entries used: *H. sapiens* (Q9UBF8-2), *M. musculus* (Q8BKC8), *D. rerio* (Q49GP3), *D. melanogaster* (Q9BKJ2), *C. elegans* (Q20077). Uniprot c10orf76 entries used: *H. sapiens* (Q5T2E6), *M. musculus* (Q6PD19), *D. rerio* (Q6PGW3), *D. melanogaster* (Q7KSU3).

### DNA Constructs and Antibodies

The following antibodies were used to examine protein localization in WT and c10orf76 knockout HAP1 cells. Mouse monoclonal antibodies included anti-GBF1 (BD Biosciences), anti-CM1 (a gift from Felix Wieland, Heidelberg University, Germany), anti-GM130 (BD Biosciences), anti-Giantin (Enzo Life Science), anti-ERGIC53 (Enzo Life Science), anti-βCOP (Sigma), anti-PI4P (Echelon). Rabbit polyclonal antibodies included anti-PI4KB (Millipore), anti-COPI α/γ (a gift from Felix Wieland), anti-TGN46 (Novus Biologicals). Conjugated goat anti-rabbit and goat anti-mouse Alexa Fluor 488, 596, or 647 (Molecular Probes) were used as secondary antibodies.

GFP-PI4KB, GFP-PI4KB RL494EA, GFP-c10orf76, and GFP-c10orf76 FLH409AAA were cloned using Gibson assembly [58] into the pEGFP-C1 vector (Clonetech). mRFP-FKBP12-PI4KB and mRFP-FKBP12-PI4KB RL494EA were generated by amplifying the mRFP-FKBP12 insert from mRFP-FKBP12-5ptpase domain [59] and replacing the N-terminal GFP in either GFP-PI4KB or GFP-PI4KB RL494EA using a single digest with NdeI. AKAP-FRB-CFP, which is used to selectively recruit FKBP12-tagged proteins to the outer mitochondrial membrane, has been described previously [60].

### Cell Culture, Transfection, and Live-Cell Confocal Microscopy of Rapamycin Recruitment

HEK293-AT1 cells, which stably express the AT1a rat Angiotensin II receptor [61], were cultured in Dulbecco’s Modified Eagle Medium (DMEM-high glucose) containing 10% (vol/vol) FBS and supplemented with a 1% solution of penicillin/streptomycin. This cell line is regularly tested for *Mycoplasma* contamination using a commercially-available detection kit (InvivoGen) and, after thawing, the cells are treated with plasmocin prophylactic (InvivoGen) at 500 µg/ml for the initial three passages (6-9 days) as well as supplemented with 5 µg/ml of plasmocin prophylactic for all subsequent passages.

For confocal microscopy, HEK293-AT1 cells (3×10^5^ cells/well) were plated on 29 mm circular glass-bottom culture dishes (#1.5; Cellvis) pre-coated with 0.01% poly-L-lysine solution (Sigma). The cells were allowed to attach overnight prior to transfection of plasmid DNAs (0.1-0.2 µg/well) using Lipofectamine 2000 (Invitrogen) and Opti-MEM (Invitrogen) according to the manufacturer’s instructions. Please note that studies using the rapamycin-inducible protein hetero-dimerization system used a 1:2:1 ratio of plasmid DNA for transfection of the FKBP12-tagged PI4KB enzyme, AKAP-FRB-CFP recruiter, and GFP-c10orf76 variant (total DNA: 0.4 µg/well). After 18-20 hr of transfection, cells were incubated in 1 mL of modified Krebs-Ringer solution (containing 120 mM NaCl, 4.7 mM KCl, 2 mM CaCl_2_, 0.7 mM MgSO_4_, 10 mM glucose, 10 mM HEPES, and adjusted to pH 7.4) and images were acquired at room temperature using a Zeiss LSM 710 laser-scanning confocal microscope (Carl Zeiss Microscopy). Rapamycin treatment of cells was carried out at a final concentration of 100 nM. Image acquisition was performed using the ZEN software system (Carl Zeiss Microscopy), while the image preparation was done using the open-source FIJI platform [62].

### Cell Culture, Transfection, and Live-Cell Confocal Microscopy of HAP1 WT and c10orf76 Knockout Cells

#### Cells and viruses

HAP1 WT cells and HAP1 c10orf76 knockout cells were obtained from Horizon Discovery. HeLa R19 cells were obtained from G. Belov (University of Maryland and Virginia-Maryland Regional College of Veterinary Medicine, US). HeLa PI4KB knockout cells were described previously [29]. HAP1 cells were cultured in IMDM (Thermo Fisher Scientific) supplemented with 10% fetal calf serum (FCS) and penicillin–streptomycin. HeLa cells were cultured in DMEM (Lonza) supplemented with 10% FCS and penicillin–streptomycin. All cells were grown at 37°C in 5% CO_2_. The following enteroviruses were used: CVA10 (strain Kowalik, obtained from the National Institute for Public Health and Environment; RIVM, The Netherlands), CVB3 (strain Nancy, obtained by transfection of the infectious clone p53CB3/T7 as described previously [63], PV1 (strain Sabin, ATCC). Virus titers were determined by end-point titration analysis and expressed as 50% tissue culture infectious dose (TCID_50_).

#### Replication rescue assay

HeLa cells were transfected with plasmids carrying WT or mutant PI4KB (RL494EA), Golgi-targeting EGFP (pEGFP-GalT) or kinase-dead PI4KB (PI4KB-KD) as a negative control. At 24 h post-transfection, the cells were infected with CVA10, CVB3, and PV1. At 8 h p.i., the infected cells were frozen, and virus titers were determined by end-point titration analysis and expressed as 50% tissue culture infectious dose (TCID_50_).

#### Immunofluorescence microscopy of WT and c10orf76 knockout HAP1 cells

HAP1 cells were grown on ibiTreat slides µ-slide 18-wells (Ibidi) one day prior to infection. Cells were fixed by submersion in a 4% paraformaldehyde solution for 15 minutes. Nuclei were stained with DAPI. Confocal imaging was performed with a Leica SpeII confocal microscope.

## Author contributions

JAM, RMH, MLJ, and JTBS expressed and purified proteins. JAM and JEB designed complex-disrupting mutations. JAM carried out pulldowns and kinase assays. JAM, MLJ, RMH and JEB carried out HDX-MS and analysis. HRL and WvE performed viral infection assays, and HRL characterized c10orf76 knockout cells. JGP and TB performed cellular c10orf76 recruitment experiments. JAM, JRPMS, TB, FJMK, and JEB designed the research. JAM and JEB wrote the manuscript with input from all authors.

### Acknowledgements

J.E.B. wishes to thank CIHR (CIHR new investigator grant and CIHR open operating grant FRN 142393) and MSFHR (scholar award 17646) for support. JAM and MLJ were supported by graduate scholarships from Natural Sciences and Engineering Research Council of Canada (NSERC). J.G.P. and T.B. are supported by the National Institutes of Health (NIH) Intramural Research Program (IRP), with additional support to J.G.P. from an NICHD Visiting Fellowship and Natural Sciences and Engineering Research Council of Canada (NSERC) Banting Postdoctoral Fellowship. Work in the lab of FJMvK is supported by research grants from the Netherlands Organization for Scientific Research (NWO-VICI-91812628, NWO-ECHO-711.017.002) and from the European Union (Horizon 2020 Marie Skłodowska-Curie ETN ‘ANTIVIRALS’, grant agreement number 642434). JRPMS is supported by a research grant from the Netherlands Organization for Scientific Research (NOW-VENI-722.012.066). The plasmid for ACBD3 and mCherry-GBF1 was a gift from Jun Sasaki and Catherine Jackson respectively. We appreciate the feedback on the manuscript pre-submission by Dr Julie Brill.

## Conflict of interest

The authors declare that they have no conflict of interest

---

**Supplemental Figure 1.**
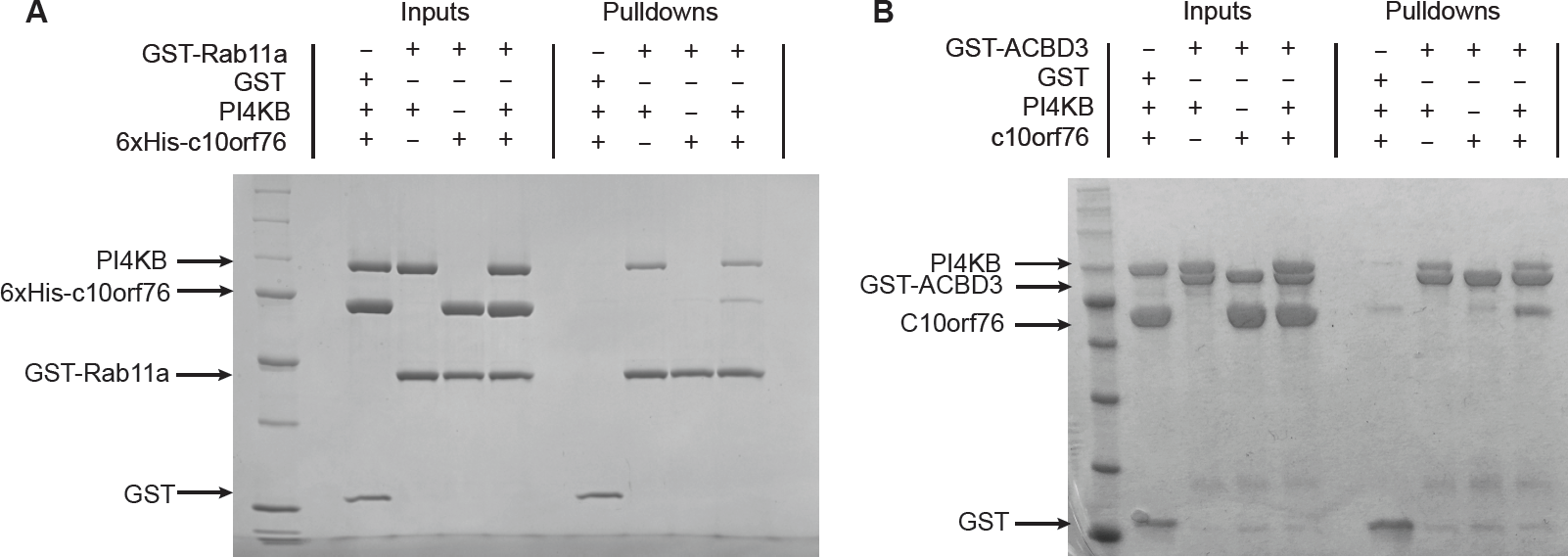
PI4KB forms ternary complexes with c10orf76, Rab11a and ACBD3. **(A)** PI4KB can form ternary complexes with Rab11a and c10orf76 *in vitro.* GST-pulldown assays were carried out using GST-Rab11a(Q70L) (6 μM) or GST alone (3 μM) as the bait, using 6xHis-c10orf76 (4 μM), PI4KB (2 μM) as the prey. **(B)** PI4KB can form ternary complexes with ACBD3 and c10orf76 *in vitro.* GST-pulldown assays were carried out using GST-ACBD3 (4 μM) or GST alone (4 μM) as the bait, and 6xHis-c10orf76 (3 μM) and PI4KB (2 μM) as the prey. Samples were washed a total of 4 times in all experiments.

**Supplemental Figure 2.**
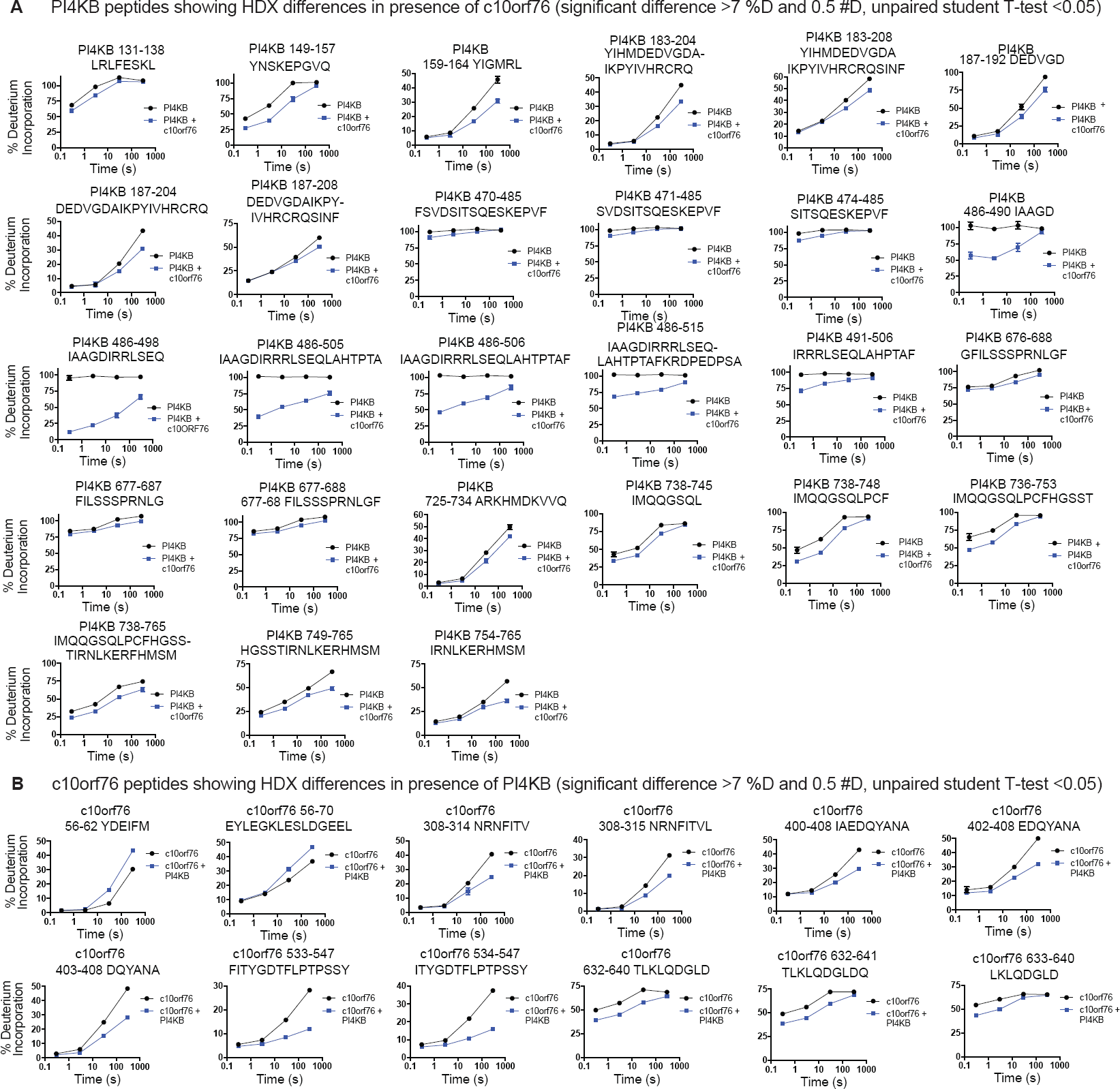
PI4KB and c10orf76 form an extended interface with spanning multiple regions. All peptides of both PI4KB **(A)** and c10orf76 **(B)** with a significant difference in H/D exchange with >7% decrease in exchange, >0.5 Da difference, and unpaired two-tailed student t-test p<0.05 at any time point (3s at 1 °C; 3s, 30s, and 300s at 23 °C).

**Supplemental Table 1.** Full statistics on all hydrogen deuterium exchange experiments according to the guidelines from the International Conference on HDX-MS.

| Data Set | PI4KB | c10orf76 | FLH409AAA<br>c10orf76 mutant |
| --- | --- | --- | --- |
| HDX reaction details | %D<sub>2</sub>O=87.4%<br>pH<sub>(read)</sub>= 7.5<br>Temp= 23°C | %D<sub>2</sub>O=87.4%<br>pH<sub>(read)</sub>= 7.5<br>Temp= 23°C | %D<sub>2</sub>O=90.5%<br>pH<sub>(read)</sub>= 7.5<br>Temp= 23° |
| HDX time course | 3s at 1°C<br>3s, 30s, 300s at<br>23°C | 3s at 1°C<br>3s, 30s, 300s at<br>23°C | 3s, 300s at 23°C |
| HDX controls | N/A | N/A | N/A |
| Back-exchange | Corrected using a<br>fully deuterated<br>(FD) sample | Corrected based on<br>%D<sub>2</sub>O | Corrected based on<br>%D<sub>2</sub>O |
| Number of peptides | 185 | 108 | 111 |
| Sequence coverage | 96.9% | 73.9% | 72.8% |
| Average peptide<br>length/ redundancy | Length = 13.8<br>Redundancy = 3.2 | Length = 12.1<br>Redundancy = 1.9 | Length = 10.7<br>Redundancy = 1.7 |
| Replicates | 3 | 3 | 3 |
| Repeatability | Average StDev =<br>1.2% | Average StDev =<br>0.6% | Average StDev =<br>1% |
| Significant<br>differences in HDX | >7% and >0.5 Da<br>and unpaired t-test <0.05 | >7% and >0.5 Da<br>and unpaired t-test<br><0.05 | >7% and >0.5 Da<br>and unpaired t-test<br><0.05 |

**Supplemental Figure 3.**
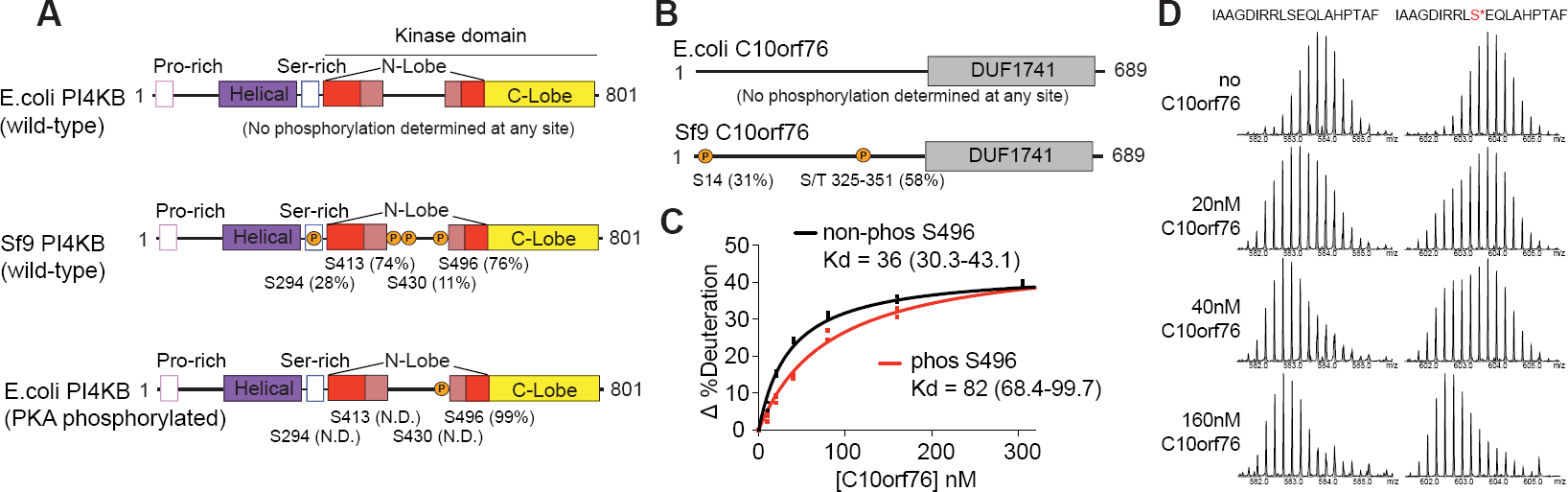
PKA phosphorylation of PI4KB Ser496 reduces affinity for c10orf76. **(A)** PKA strongly phosphorylates Ser496 over Ser294 and Ser413 sites. Relative abundance of phosphorylated PI4KB at Ser294, Ser413 and Ser496 sites were calculated using the relative intensity (total area) of the phosphorylated vs non-phosphorylated peptides (peptides 290-303, 290-312, 395-425, 430-441, 486-505 and 486-506) for PI4KB expressed in *Sf9*, expressed in *E. coli*, and expressed in *E. coli* and treated with PKA. N.D. indicates no phosphorylation was identified. N.D. indicates no phosphorylation was identified. **(B)** c10orf76 contains two phosphorylation sites when produced in *Sf9.* Relative abundance of S14 was calculated using the relative intensity (total area) of the phosphorylated vs non-phosphorylated peptide (10-22). Relative abundance of the second phosphorylation site in the 325-351 region was calculated using the relative intensity (total area) average over three phosphorylated vs non-phosphorylated peptides; the definitive Ser/Thr phosphorylation residue could not be determined. **(C)** Phosphorylation of Ser496 reduces PI4KB affinity for c10orf76. Deuterium incorporation of the PI4KB kinase linker region peptide 488-508 (20 nM) at a single time point (5 seconds of D_2_O exposure at 23°C) was monitored in the presence of increasing concentrations of c10orf76 (0-320 nM c10orf76). K_d_ values were generated using a one binding site, nonlinear regression (curve fit), and are shown with 95% confidence intervals. Error bars represent standard deviation (n=3). **(D)** Raw deuterium incorporation data for PI4KB peptide 488-508 used to generate panel C. The deuterium incorporation for the phosphorylated and non-phosphorylated variants of PI4KB are shown in the presence of different concentrations of c10orf76.

**Supplemental figure 4.**
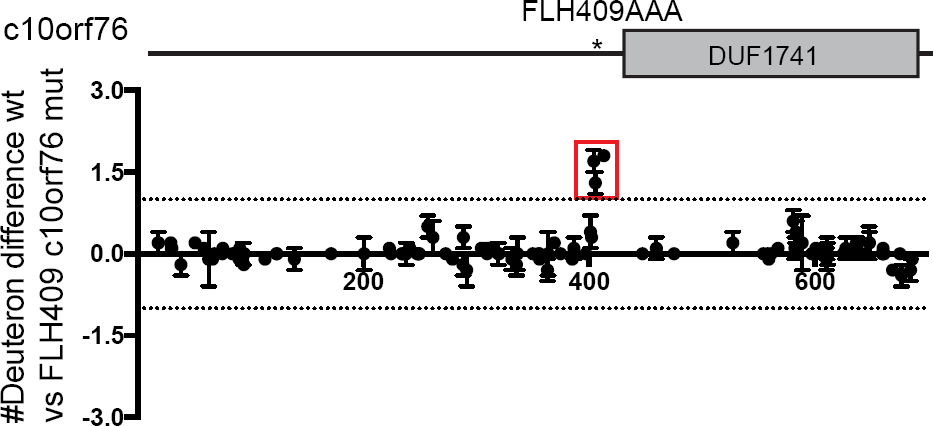
The FLH409AAA c10orf76 mutant maintains similar overall secondary structure to wild-type with a destabilization at the mutation site. Differences in the changes in the deuterium incorporation of wild type and FLH409AAA mutant c10orf76. H/D exchange reactions of c10orf76 (400 nM) were carried out for 3s and 300s, and the average difference in number of deuterons incorporated between wild-type and FLH409AAA c10orf76 (400 nM) was graphed. Error bars represent standard deviation (n=3).

